# StackGlyEmbed: Prediction of N-linked Glycosylation sites using protein language models

**DOI:** 10.1101/2025.02.12.637996

**Authors:** Md Muhaiminul Islam Nafi, M Saifur Rahman

**Affiliations:** Department of CSE, BUET, Dhaka 1000, Bangladesh

**Author notes:** **Md Muhaiminul Islam Nafi** Currently an undergraduate student at the Bangladesh University of Engineering and Technology (BUET) in the Department of CSE. **M Saifur Rahman** Currently a professor at the Bangladesh University of Engineering and Technology (BUET) in the Department of CSE.

**Keywords:** Embeddings, ProtT5-XL-U50, per-residue, window, Glycosylation, Stacking ensemble, SHAP

## Abstract

N-linked glycosylation is one of the most basic post-translational modifications (PTMs) where oligosaccharides covalently bond with Asparagine (N). These are found in the conserved regions like N-X-S or N-X-T where X can be any residue except Proline (P). Prediction of N-linked glycosylation sites has great importance as these PTMs play a vital role in many biological processes and functionalities. Experimental methods, such as mass spectrometry, for detecting N-linked glycosylation sites are very expensive. Therefore, prediction of N-linked glycosylation sites has become an important research field. In this work, we propose StackGlyEmbed, a stacking ensemble machine learning model, to computationally predict N-linked glycosylation sites. We have explored embeddings from several protein language models and built the stacking ensemble using SVM, XGB and KNN learners in the base layer, with a second SVM model in the meta layer. StackGlyEmbed achieves 98.2% sensitivity, 92.5% balanced accuracy, 89.1% F1-score and 82.6% MCC in independent testing, outperforming the existing SOTA methods. StackGlyEmbed is freely available at https://github.com/nafcoder/StackGlyEmbed.

## 1 Introduction

Post-translational modifications (PTMs) refer to side chain modification of amino acids in proteins after their biosynthesis [1]. N-linked glycosylation is one of the most basic PTMs where Asparagine (N) covalently bonds with oligosaccharides, thus producing N-glycans. This PTM appears in highly conserved areas like N-X-S or N-X-T [2] [3]. Here, in N-X-[S/T], the 1st position will be Asparagine (N), the 2nd position can be any residue except Proline (P) and the 3rd position can be Serine (S) or Threonine (T). N-linked glycosylation is necessary for the structure, stability and functionality of glycoproteins in both prokaryotic and eukaryotic organisms. It plays a crucial role in protein folding and stability, biological functions, immune responses, cell-to-cell communication and clinical diagnosis. N-X-[S/T] does not guarantee that the site is N-linked glycosylated [4] [5] [6] [7]. Other factors like distance to the next glycosylated site, and neighboring residues of the potential sequon can also have an effect on its being glycosylated or not. For N-linked glycosylation to take place, having N-X-[S/T] sequon is therefore necessary but not sufficient [2] [4] [7] [8].

Methods like radioactive chemical labeling [9] and mass spectrometry [10] [11] are utilized for the identification of N-linked glycosylation. As these methods are very expensive, computationally predictive methods have been created to predict N-linked glycosylation over the years. Gupta and Brunak [12] crafted glycosylation site prediction methods for N-linked, O-linked GalNAc (mucin-type) and O-*β*-linked GlcNAc (intracellular/nuclear) glycosylations using Artificial Neural Networks (ANN) that examined correlations in the local sequence context and surface accessibility. Subsequently, Gupta et al. [13] designed NetNGlyc, an ANN-based method, that predicted N-glycosylation sites in protein sequences. Chauhan et al. [14] developed GlycoPP which is an SVM-based prediction model trained on O- and/or N-glycosylated proteins from six different archaeal and bacterial phyla. The features were different sequential properties. Chuang et al. [15] proposed NGlycPred that utilized both structural and residue pattern information using the Random Forest algorithm on a set of N-linked glycosylated protein structures. Chauhan et al. [16] created GlycoEP, a Support Vector Machine (SVM) classifier model, which predicted N-linked, C-linked and O-linked glycosites in eukaryotic glycoproteins using two larger datasets, namely, standard and advanced datasets.

Li et al. [17] constructed GlycoMine using the Random Forest (RF) algorithm for computer-based identification of C-linked, N-linked, and O-linked glycosylation sites belonging to the human proteome. Pitti et al. [18] built N-GlyDE, a two-stage prediction tool, that predicted N-linked glycosylation sites in human proteins. The first stage used a protein similarity voting algorithm and the second stage used SVM. Taherzadeh et al. [19] proposed SPRINT-Gly that used Deep Neural Network (DNN) for predicting N-linked glycosylation sites for humans and mice. Pugalenthi et al. [20] developed Nglyc, a Random Forest Method, for the prediction of N-Glycosylation sites in proteins related to humans and mice. Chien et al. [21] manufactured N-GlycoGo, a predicting tool, using XGBoost for predicting N-Glycosylation sites on humans and mouse glycoproteins. Pakhrin et al. [22] designed DeepNGlyPred, a deep learning-based approach, that trained and tested on the human proteome datasets (N-GlyDE and N-GlycositeAtlas). Pakhrin et al. [23] developed LMNglyPred, a deep learning-based approach, to predict N-linked glycosylated sites in human proteins (N-GlyDE and N-GlycositeAtlas). They used embeddings from a pre-trained protein language model (ProtT5-XL-U50 [24]) as features. Hou et al. [25] designed EMNGly, an SVM-based model, for predicting N-linked glycosylation sites from a variety of species, including humans, mice, rats, and yeast.

As discussed earlier, for the N-linked glycosylation to happen, it is a necessary (but not sufficient) condition that the target site must form an N-X-[S/T] sequon. Therefore, to train an N-linked glycosylation predictor, the dataset should consist only of N-X-[S/T] sequons, some of which are positive sites, and some are not. However, many earlier works used datasets consisting of sequons that did not adhere to the above motif. Recent works, such as NetNGlyc, N-GlyDE, DeepNGlyPred, LMNglyPred, and EMNGly, have on the other hand only focused on datasets comprising N-X-[S/T] sequons and our work follows this approach as well.

A majority of these existing predictors related to N-linked glycosylation depend on input features that have been manually crafted. This dependency creates a bias towards certain characteristics and hinders the discovery of underlying representations of important but unfamiliar features. In our study, we used embedding features, obtained from various protein language models (PLMs), such as ProteinBERT [26], ESM-2 [27], and ProtT5-XL-U50 [24] embeddings, that are known for their wide range of successful usage in multiple protein attribute prediction tasks [28– 31]. In addition to per-residue embedding, we have also utilized window representations that consider the neighboring residues for an aggregate embedding. We have also experimented with the physicochemical properties of the residues as features. To the best of our knowledge, ours is the first study to combine ProteinBERT, ESM-2, and ProtT5-XL-U50 embeddings to address the challenges of N-linked glycosylation prediction. Additionally, existing studies often overlook the importance of selecting specific feature groups for prediction. To address this, we performed Incremental Feature Selection (IFS) on various feature groups to identify the optimal feature set for our final model. We experimented with two different datasets using 10 different ML classifiers and different feature groups. Per-residue as well as window features were examined. Interestingly, feature selection using IFS led to the same set of final features in both datasets. We utilized Incremental Mutual Info (IMI) for selecting the base classifiers for our final model. Then, we did a 10-Fold CV to choose the meta-classifier for our ensemble model. The learners/classifiers were further fine-tuned using grid search. Finally, we did a SHAP (SHapley Additive exPlanations) [32] analysis on the selected feature set which proved the significance of base learner probability (BLP) outputs in the stacking ensemble.

The key contributions of this paper can be summarized as follows.

- We have proposed StackGlyEmbed, a stacking ensemble method for N-linked glycosylation site prediction. StackGlyEmbed outperforms existing SOTA methods in independent datasets, achieving 98.2% sensitivity, 92.5% balanced accuracy, 89.1% F1-score and 82.6% MCC.
- StackGlyEmbed utilizes multiple protein language models, such as, ProtT5-XL-U50, ProteinBERT and ESM2. To the best of our knowledge, ours is the first study that combines embeddings from multiple protein language models to predict N-linked glycosylation. We have analyzed the performance of each of the feature groups and selected our model feature set using IFS.
- In addition to using per-residue embedding, we have also used aggregate embedding across a window around the target site for the prediction task. Notably, the *windowing* approach has not been explored enough in the recent literature on N-linked glycosylation prediction.
- We have analyzed the selected feature groups using SHAP [32] and demonstrated the benefit of the stacking ensemble.
- We have made our proposed model freely available at https://github.com/nafcoder/StackGlyEmbed.

The rest of the sections are organized as follows. Section 2 outlines the materials and methods used in this paper. Section 3 presents the results of different experiments, performance evaluation, and comparative analysis of our approach. Section 4 explores the implications of our findings in the context of N-linked glycosylation prediction and concludes the paper with potential future directions.

## 2 Materials and Methods

This section covers the datasets we have used, feature extraction and selection methods, our model framework, etc.

### 2.1 Dataset

In this subsection, two datasets: N-GlyDE and N-GlycositeAtlas are described. The dataset summary is provided in Table 1.

**Table 1:** Dataset summary.

| Dataset | Positive sites | Negative sites | Total sites | Proteins | Total proteins |
| --- | --- | --- | --- | --- | --- |
| N-GlyDE-TS | 2049 | 1030 | 3079 | 832 | 918 |
| N-GlyDE-IT | 166 | 280 | 446 | 86 |  |
| N-GlycositeAtlas-TS | 8401 | 15857 | 24258 | 9245 | 10200 |
| N-GlycositeAtlas-IT | 830 | 1648 | 2478 | 955 |  |

#### 2.1.1 N-GlyDE dataset

This dataset was taken from N-GlyDE [18]. The authors used Uniprot [33] (version 201608) for preparing their datasets. From Uniprot, they collected 1068 experimentally validated N-linked human glycoproteins, and 11767 non-glycoproteins. After some post-processing, and ensuring a 30% sequence identity threshold using CD-HIT [34], 6281 proteins remained. From these, 86 proteins (comprising 53 glycoproteins and 33 non-glycoproteins), were randomly selected to prepare the independent test set. Finally, 447 N-X-[S/T] sequons were compiled from these proteins, of which 167 were glycosites (positive sites) and 280 were non-glycosites (negative sites). For training purpose, they created two datasets, referred to as the *first-stage dataset* and *second-stage dataset*. In our work, we have used the latter one as the training set. To prepare this dataset (*second-stage dataset*), the authors relaxed the sequence identity threshold to 60% and got 832 glycoproteins with a total of 3080 sequons. 2050 of them were glycosites and 1030 were non-glycosites. They also used CD-HIT to ensure that the protein sequence identity between the independent test set and the training set remains at most 30%. For a more detailed review of the data preparation protocol, the reader is referred to [18]. From the original datasets, we had to remove two sequons due to the corresponding protein’s primary sequence having been updated or shortened in Uniprot. In our study, the final training set (N-GlyDE-TS) thus has 2049 positive sites and 1030 negative sites, while the final independent test set (N-GlyDE-IT) has 166 positive sites and 280 negative sites (see Table 1). We further divided N-GlyDE-TS into two parts: 60% (N-GlyDE-TS-60) for training the base layer classifiers and 40% (N-GlyDE-TS-40) for validation of base layer, and training of the meta layer classifier. N-GlyDE-TS-60 contains 1229 positive and 618 negative sites, while N-GlyDE-TS-40 consists of 820 positive and 412 negative sites.

### 2.1.2 N-GlycositeAtlas dataset

This dataset was taken from LMNglyPred [23], where the authors took 7204 human glycoproteins from N-GlycositeAtlas [35]. From these proteins, 9235 N-linked glycosites were obtained that were confined to N-X-[S/T] sequons. On the other hand, to construct the negative set, the authors extracted 7875 human glycoproteins from the DeepLoc-2.0 database [36]. 30% sequence identity cutoff was ensured using CD-HIT [37]. Finally, 17508 non-glycosites were obtained, confined to N-X-[S/T] sequons. From these 9235 glycosites and 17508 non-glycosites, the authors then formed a training set and an independent test set. 8405 glycosites and 15860 non-glycosites belonged to the training set while the independent test set comprised 830 glycosites and 1648 non-glycosites. For a more detailed review of the data preparation protocol, the reader is referred to [23]. From the original training set, we had to remove seven sequons due to the corresponding protein’s sequence having been updated or shortened in Uniprot, thus reducing its composition to 8401 positive and 15857 negative sites. The composition of the training set (N-GlycositeAtlas-TS) and independent test set (N-GlycositeAtlas-IT) is summarized in Table 1. Similar to the training and validation split performed in the N-GlyDE-TS, we further split N-GlycositeAtlas-TS into N-GlycositeAtlas-TS-60 (5040 positive and 9514 negative sites) and N-GlycositeAtlas-TS-40 (3361 positive and 6343 negative sites) sets.

### 2.2 Feature Extraction

In this work, we have utilized physicochemical properties and embeddings from protein language models as features for the prediction problem. These are described below.

#### 2.2.1 Physicochemical properties

We used seven different physicochemical parameters (sheet probability, steric parameter, hydrophobicity, isoelectric point, normalized Van der Waals volume, helix probability and polarizability) for each of the 20 standard amino acids. We took a window of 31 residues, centered at the target site, and averaged their physicochemical parameter values. The feature group, named Physico-Window, thus had a size of 7.

#### 2.2.2 Embedding from ProtT5-XL-U50

ProtTrans [24] has explored two auto-regressive models (Transformer-XL, XLNet) and four auto-encoder models (BERT, Albert, Electra, T5) on 393 billion amino acids of UniRef and BFD data. Among the available ProTrans models, ProtT5-XL-U50 has the highest score in CASP12, TS115, CB513 and DeepLoc for various predictions. So, we used ProtT5-XL-U50. We computed the per-residue embeddings of size 1024 and labeled them as ProtT5-XL-U50-Per-Residue. We also took a window of 31 residues, centered at the target site, and averaged their per-residue embeddings. This feature group is named ProtT5-XL-U50-Window.

#### 2.2.3 Embedding from ESM2

ESM2 [27] is a transformer-based protein language model that is trained using masked language modeling over millions of diverse natural proteins across evolution. It is trained for getting the inter-connected sequence patterns and their structural implications. It generates per-residue embeddings that are later used for unsupervised selfattention map contact predictions. This is then used by ESMFold [27] for predicting protein structures. We used esm2 t33 650M UR50D for our computations. We generated per-residue embeddings of size 1280 (ESM-2-Per-Residue) as well as embeddings averaged across a window of 31 residues (ESM-2-Window).

#### 2.2.4 Embedding from ProteinBERT

ProteinBERT [26], a deep language model, was designed to capture local and global representations of proteins naturally. Its architecture consists of local and global representations that were pre-trained on around 106M proteins. We took the protein sequence and computed the global representation of it. It had an embedding size of 512.

### 2.3 Feature selection

For the feature selection process, we used Incremental Feature Selection (IFS), with SVM as the classifier. Firstly, we measured the F1-score of SVM models trained on each feature group individually and opted for the best feature group. In the next round, we augmented this feature group with each remaining feature group individually. We trained SVM models again, now on each of these feature group pairs, and measured their F1-scores. The pair that resulted in the best F1-score was kept for subsequent rounds, and we continued with this process until we reached the feature set containing all of the feature groups. Finally, among all the feature group sets thus explored, the one with the best F1-score was chosen for the final model. Accordingly, ProteinBERT, ESM-2-Window and ProtT5-XL-U50-Per-Residue embeddings were selected as the optimal feature set. The selected feature set is shown in Figure 1.

**Figure 1.**
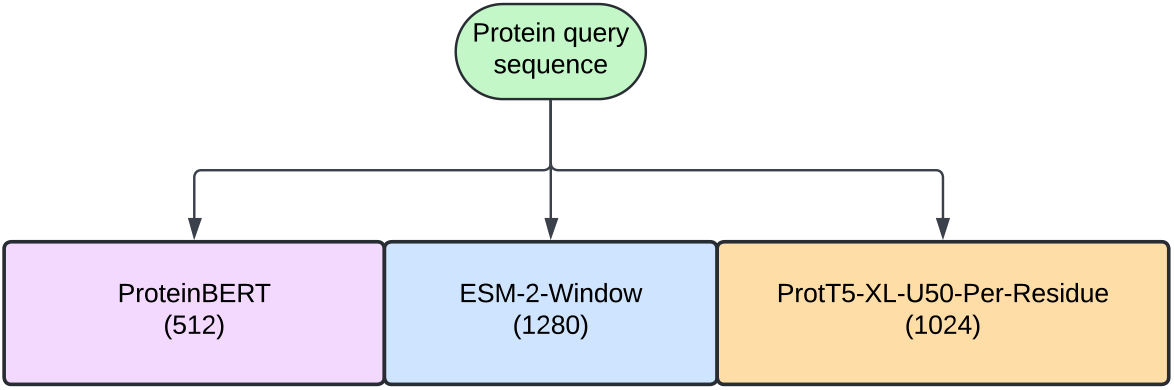
Selected feature set

### 2.4 Performance evaluation

We have used various well-established metrics [38] [39] [40] [41] for measuring the performance of our model. The metrics we used are balanced accuracy (BACC), accuracy (ACC), specificity (SP), sensitivity (SN), F1-score (F1), precision (PREC), Matthew’s Correlation Coefficient (MCC), area under precision-recall curve (AUPR) and area under receiver operating characteristic curve (AUROC or AUC).

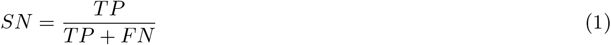

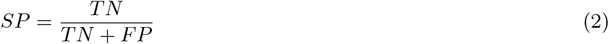

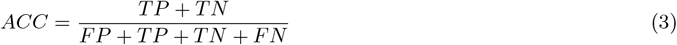

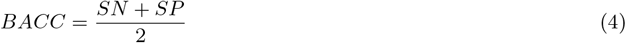

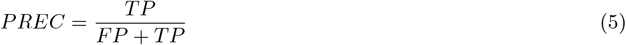

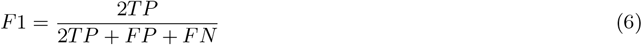

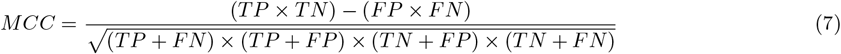

Here, TP, FP, TN, FN, TPR and FPR represent True Positives, False Positives, True Negatives, False Negatives, True Positive Rate and False Positive Rate respectively.

### 2.5 Summary of the framework

We constructed a stacking ensemble with two phases of learners, referred to as *base learners* in the base layer and a single *meta learner* in the meta layer, respectively. Two separate ensembles were trained on the two datasets, N-GlyDE and N-GlycositeAtlas, following an identical methodology. Therefore, we describe the process for the N-GlyDE dataset only. Notably, the data was pre-processed using Yeo–Johnson transformation [42] to make it more Gaussian-like. We used the N-GlyDE-TS-60 set to train the base learners, while N-GlyDE-TS-40 was used to train the meta learner. The data was featurized using the optimal feature set determined by the IFS method described earlier. When training the meta layer, the feature vectors for N-GlyDE-TS-40 were augmented using the probabilities obtained from the base layer, referred to as the Base Learner Probabilities (BLP) vector.

The selection of base learners was made using the Incremental Mutual Information (IMI) approach that utilized Mutual Information (MI) [43] between learner outputs and the actual labels. IMI works similarly to the IFS method. The only differences are that IMI is searching an optimal set of learners (instead of an optimal feature set) and the objective function here is mutual information (instead of F1-score). We analyzed 10 classifiers in total which are: Support Vector Machine (SVM), Extreme Gradient Boosting (XGB), Extra Tree (ET), Multi-layer Perceptron (MLP), Partial Least Square Regression (PLS), Logistic Regression (LR), Naive Bayes (NB), Random Forest (RF), Decision Tree (DT) and *K*-nearest Neighbor (KNN). IMI resulted in the selection of SVM, XGB and KNN as the base learners. On the other hand, SVM was chosen as the meta learner since it was the best individual classifier in terms of predictive performance (see Results). The overall framework for training and inference is shown in Figure 2.

**Figure 2.**
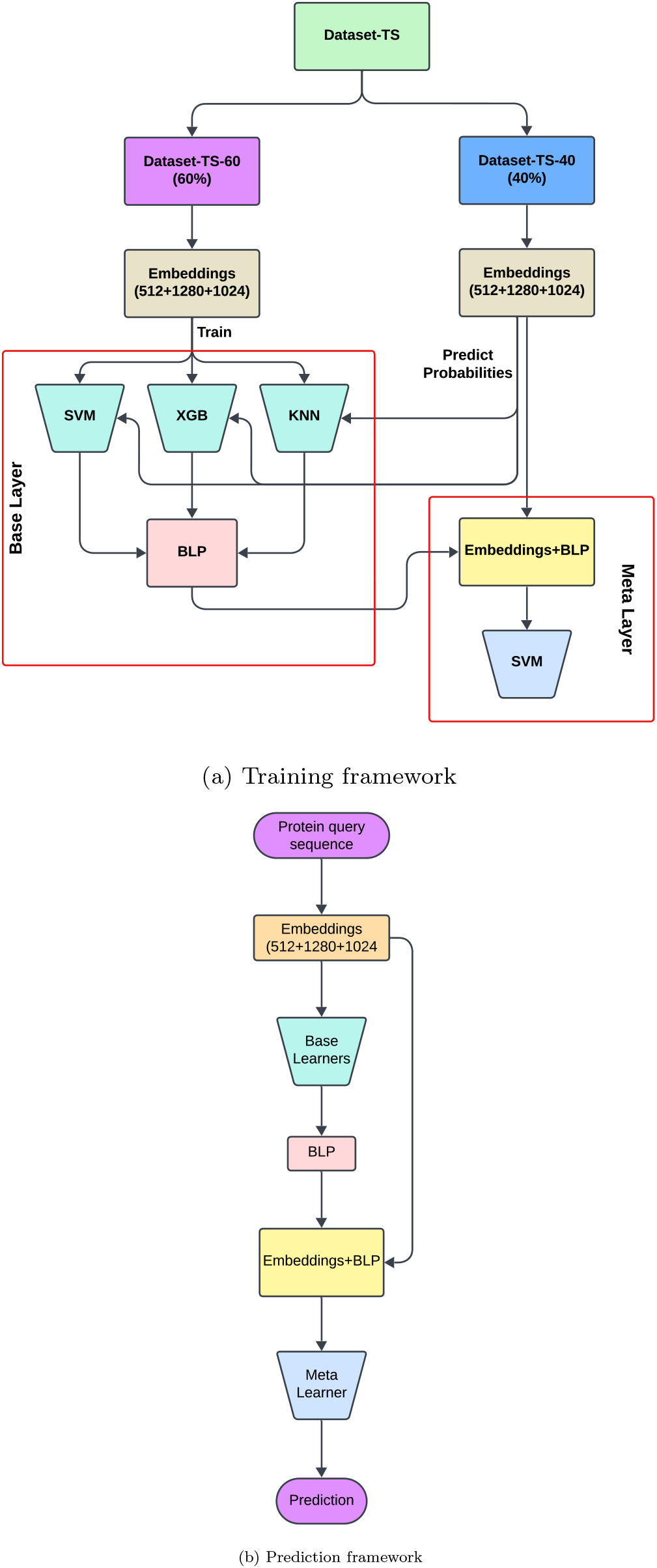
Training and prediction framework

### 2.6 Resolving data imbalance

There is a notable imbalance in the datasets used in this research. For N-GlyDE the positive sites are almost double the number of the negative sites while it is the other way around for N-GlycositeAtlas. Inspired by [44], we followed an ensemble random undersampling strategy for the N-GlyDE-TS-60 and N-GlycositeAtlas-TS-60 datasets to train the base learners. We describe the process for N-GlyDE-TS-60 only. Ten subsampled datasets were created from it including all sites of the minority class and varying percentages of sites from the majority class - three samples contained 70% of the majority class, four contained 50%, and three contained 30%. The three base learners (SVM, XGB and KNN) were trained on these 10 datasets, thus rendering thirty trained models in the base layer. For the meta learner training, on the other hand, the N-GlyDE-TS-40 and N-GlycositeAtlas-TS-40 datasets were balanced using random undersampling.

## 3 Results

### 3.1 ProteinBERT, ESM-2-Window and ProtT5-XL-U50-Per-Residue constitute the optimal feature set

As mentioned in Section 2.3, we used IFS with SVM for feature selection. The choice of SVM here was not arbitrary. Rather, each of the 10 ML classifiers were trained separately on each of the 6 feature groups. For each classifier, we recorded various performance metrics, averaged across the feature groups, in Tables 2 and 3, for the two different training datasets respectively. In both cases, SVM achieved the best performance based on F1, MCC, AUROC and AUPR scores, which justified its incorporation with the IFS method for feature selection.

**Table 2:** 10-Fold CV performance of all 10 classifiers, averaged over all 6 feature groups, trained on the N-GlycositeAtlas-TS set that was balanced using random undersampling.

| Classifier | SN | SP | BACC | ACC | PREC | F1 | MCC | AUROC | AUPR |
| --- | --- | --- | --- | --- | --- | --- | --- | --- | --- |
| SVM | **0.812** | 0.669 | **0.74** | **0.74** | 0.715 | **0.759** | **0.485** | **0.805** | **0.776** |
| XGB | 0.736 | 0.687 | 0.712 | 0.712 | 0.703 | 0.719 | 0.424 | 0.777 | 0.747 |
| ET | 0.728 | 0.683 | 0.705 | 0.705 | 0.697 | 0.712 | 0.411 | 0.76 | 0.709 |
| MLP | 0.739 | **0.71** | 0.724 | 0.724 | **0.724** | 0.73 | 0.449 | 0.793 | 0.767 |
| PLS | 0.702 | 0.614 | 0.658 | 0.658 | 0.644 | 0.67 | 0.32 | 0.716 | 0.672 |
| LR | 0.722 | 0.674 | 0.698 | 0.698 | 0.688 | 0.704 | 0.397 | 0.767 | 0.741 |
| NB | 0.66 | 0.576 | 0.618 | 0.618 | 0.612 | 0.633 | 0.239 | 0.663 | 0.623 |
| RF | 0.723 | 0.686 | 0.705 | 0.705 | 0.697 | 0.71 | 0.41 | 0.766 | 0.724 |
| DT | 0.621 | 0.637 | 0.629 | 0.629 | 0.633 | 0.626 | 0.258 | 0.633 | 0.596 |
| KNN | 0.695 | 0.657 | 0.676 | 0.676 | 0.672 | 0.682 | 0.353 | 0.734 | 0.692 |

**Table 3:** 10-Fold CV performance of all 10 classifiers, averaged over all 6 feature groups, trained on the N-GlyDE-TS set that was balanced using random undersampling.

| Classifier | SN | SP | BACC | ACC | PREC | F1 | MCC | AUROC | AUPR |
| --- | --- | --- | --- | --- | --- | --- | --- | --- | --- |
| SVM | **0.858** | 0.689 | **0.773** | **0.773** | 0.736 | **0.791** | **0.557** | **0.831** | **0.804** |
| XGB | 0.797 | 0.707 | 0.752 | 0.752 | 0.73 | 0.761 | 0.509 | 0.807 | 0.779 |
| ET | 0.799 | 0.687 | 0.743 | 0.743 | 0.719 | 0.754 | 0.494 | 0.797 | 0.763 |
| MLP | 0.775 | **0.739** | 0.757 | 0.757 | **0.752** | 0.762 | 0.516 | 0.818 | 0.79 |
| PLS | 0.798 | 0.646 | 0.722 | 0.722 | 0.689 | 0.737 | 0.455 | 0.773 | 0.743 |
| LR | 0.708 | 0.731 | 0.719 | 0.719 | 0.725 | 0.716 | 0.44 | 0.774 | 0.752 |
| NB | 0.824 | 0.615 | 0.72 | 0.72 | 0.684 | 0.746 | 0.452 | 0.757 | 0.7 |
| RF | 0.799 | 0.685 | 0.742 | 0.742 | 0.717 | 0.754 | 0.491 | 0.801 | 0.77 |
| DT | 0.634 | 0.676 | 0.655 | 0.655 | 0.665 | 0.647 | 0.312 | 0.659 | 0.617 |
| KNN | 0.772 | 0.71 | 0.741 | 0.741 | 0.73 | 0.747 | 0.488 | 0.795 | 0.747 |

The results of IFS are recorded in Tables 4 (for N-GlycositeAtlas) and 5 (for N-GlyDE). In both cases, *{*ProteinBERT, ESM-2-Window, ProtT5-XL-U50-Per-Residue*}* feature set resulted in the highest F1-score. Therefore, it was selected as the optimal feature set. The length of the final feature vector is 2816, with each individual feature group’s feature count shown in Figure 1. For ease of subsequent reference, we have named this feature combination as FC-1. When augmented with BLP, we refer to it as FC-2 (see Table 11).

**Table 4:** 10-Fold CV performance of SVM classifier, trained on N-GlycositeAtlas-TS, using IFS. Data was balanced using random undersampling. The feature combination with the highest F1-score is chosen (shown in boldface).

| Feature set | SN | SP | BACC | ACC | PREC | F1 | MCC | AUROC | AUPR |
| --- | --- | --- | --- | --- | --- | --- | --- | --- | --- |
| Physico-Window | 0.721 | 0.436 | 0.579 | 0.579 | 0.561 | 0.631 | 0.164 | 0.607 | 0.579 |
| ProtT5-XL-U50-Per-Residue | 0.839 | 0.686 | 0.763 | 0.763 | 0.728 | 0.779 | 0.532 | 0.834 | 0.802 |
| ProtT5-XL-U50-Window | 0.848 | 0.703 | 0.775 | 0.775 | 0.741 | 0.79 | 0.556 | 0.842 | 0.806 |
| ESM-2-Per-Residue | 0.806 | 0.682 | 0.744 | 0.744 | 0.717 | 0.759 | 0.492 | 0.814 | 0.78 |
| ESM-2-Window | 0.832 | 0.715 | 0.774 | 0.774 | 0.745 | 0.786 | 0.552 | 0.848 | 0.819 |
| ProteinBERT | 0.825 | 0.789 | 0.807 | 0.807 | 0.797 | 0.811 | 0.615 | 0.884 | 0.87 |
| ProteinBERT, Physico-Window | 0.827 | 0.786 | 0.807 | 0.807 | 0.795 | 0.81 | 0.614 | 0.886 | 0.874 |
| ProteinBERT, ProtT5-XL-U50-Per-Residue | 0.854 | 0.742 | 0.798 | 0.798 | 0.768 | 0.809 | 0.6 | 0.871 | 0.851 |
| ProteinBERT, ProtT5-XL-U50-Window | 0.856 | 0.743 | 0.799 | 0.799 | 0.769 | 0.81 | 0.603 | 0.871 | 0.846 |
| ProteinBERT, ESM-2-Per-Residue | 0.833 | 0.73 | 0.782 | 0.782 | 0.756 | 0.792 | 0.567 | 0.854 | 0.83 |
| ProteinBERT, ESM-2-Window | 0.851 | 0.753 | 0.802 | 0.802 | 0.775 | 0.811 | 0.607 | 0.875 | 0.855 |
| ProteinBERT, ESM-2-Window, Physico-Window | 0.851 | 0.753 | 0.802 | 0.802 | 0.775 | 0.811 | 0.607 | 0.875 | 0.855 |
| **ProteinBERT, ESM-2-Window, ProtT5-XL-U50-Per-Residue** | **0.857** | **0.749** | **0.803** | **0.803** | **0.774** | **0.813** | **0.61** | **0.877** | **0.858** |
| ProteinBERT, ESM-2-Window, ProtT5-XL-U50-Window | 0.86 | 0.741 | 0.8 | 0.8 | 0.769 | 0.811 | 0.605 | 0.873 | 0.849 |
| ProteinBERT, ESM-2-Window, ESM-2-Per-Residue | 0.839 | 0.739 | 0.789 | 0.789 | 0.763 | 0.799 | 0.581 | 0.863 | 0.842 |
| ProteinBERT, ESM-2-Window, ProtT5-XL-U50-Per-Residue, Physico-Window | 0.856 | 0.749 | 0.803 | 0.803 | 0.774 | 0.813 | 0.609 | 0.877 | 0.858 |
| ProteinBERT, ESM-2-Window, ProtT5-XL-U50-Per-Residue, ProtT5-XL-U50-Window | 0.857 | 0.744 | 0.8 | 0.8 | 0.77 | 0.811 | 0.605 | 0.875 | 0.855 |
| ProteinBERT, ESM-2-Window, ProtT5-XL-U50-Per-Residue, ESM-2-Per-Residue | 0.849 | 0.738 | 0.794 | 0.794 | 0.765 | 0.804 | 0.591 | 0.867 | 0.844 |
| ProteinBERT, ESM-2-Window, ProtT5-XL-U50-Per-Residue, Physico-Window, ProtT5-XL-U50-Window | 0.857 | 0.744 | 0.8 | 0.8 | 0.77 | 0.811 | 0.604 | 0.875 | 0.855 |
| ProteinBERT, ESM-2-Window, ProtT5-XL-U50-Per-Residue, Physico-Window, ESM-2-Per-Residue | 0.848 | 0.739 | 0.793 | 0.793 | 0.765 | 0.804 | 0.591 | 0.867 | 0.845 |
| ProteinBERT, ESM-2-Window, ProtT5-XL-U50-Per-Residue, Physico-Window, ProtT5-XL-U50-Window, ESM-2-Per-Residue | 0.853 | 0.737 | 0.795 | 0.795 | 0.764 | 0.806 | 0.594 | 0.869 | 0.847 |

**Table 5:**
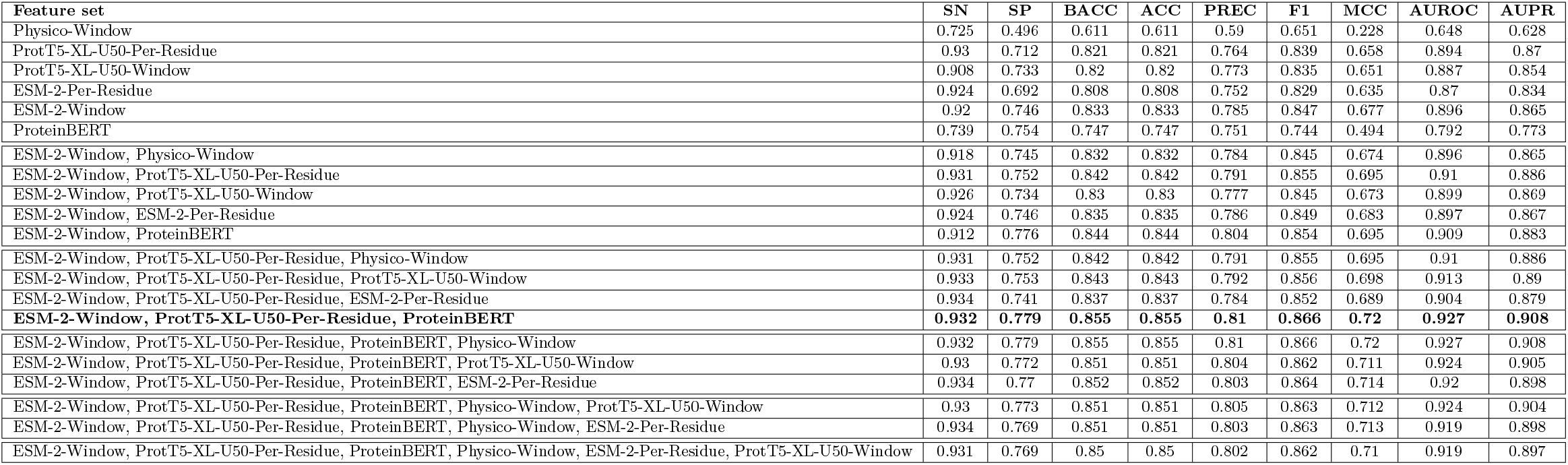
10-Fold CV performance of SVM classifier, trained on N-GlyDE-TS, using IFS. Data was balanced using random undersampling. The feature combination with the highest F1-score is chosen. In case of a tie, the combination with a lesser number of feature groups is selected (shown in boldface).

### 3.2 Choice of learners in base and meta layers

The set of classifiers having the highest MI score on each round using Incremental Mutual Information (IMI) are shown in Table 6 (for N-GlycositeAtlas) and 7 (for N-GlyDE). Full tables with performance metrics are given in Supplementary. In both cases, the combination of SVM, XGB, and KNN produced the highest MI score. Therefore, these classifiers were chosen as the base learners.

**Table 6:** Highest MI scores of the learner combinations in each round. The learners were trained on the randomly undersampled N-GlycositeAtlas-TS-60 set and MI scores were measured on the N-GlycositeAtlas-TS-40 set.

| Learner Combination | MI |
| --- | --- |
| SVM | 0.2241 |
| SVM, XGB | 0.2345 |
| **SVM, XGB, KNN** | **0.239** |
| SVM, XGB, KNN, PLS | 0.2361 |
| SVM, XGB, KNN, PLS, MLP | 0.2358 |
| SVM, XGB, KNN, PLS, MLP, RF | 0.2376 |
| SVM, XGB, KNN, PLS, MLP, RF, LR | 0.2317 |
| SVM, XGB, KNN, PLS, MLP, RF, LR, ET | 0.2239 |
| SVM, XGB, KNN, PLS, MLP, RF, LR, ET, NB | 0.2178 |
| SVM, XGB, KNN, PLS, MLP, RF, LR, ET, NB, DT | 0.2066 |

**Table 7:** Highest MI scores of the learner combinations in each round. The learners were trained on the randomly undersampled N-GlyDE-TS-60 set and MI scores were measured on the N-GlyDE-TS-40 set.

| Learner Combination | MI |
| --- | --- |
| XGB | 0.3316 |
| XGB, KNN | 0.3425 |
| **XGB, KNN, SVM** | **0.3664** |
| XGB, KNN, SVM, NB | 0.3572 |
| XGB, KNN, SVM, NB, ET | 0.3518 |
| XGB, KNN, SVM, NB, ET, MLP | 0.3498 |
| XGB, KNN, SVM, NB, ET, MLP, LR | 0.3522 |
| XGB, KNN, SVM, NB, ET, MLP, LR, DT | 0.353 |
| XGB, KNN, SVM, NB, ET, MLP, LR, DT, PLS | 0.3422 |
| XGB, KNN, SVM, NB, ET, MLP, LR, DT, RF, PLS | 0.3299 |

After settling on the base learners, we experimented with the potential meta learners as follows. We measured the 10-Fold CV performance of 10 ML models on the randomly undersampled N-GlycositeAtlas-TS-40 set and N-GlyDE-TS-40 set in Table 8 and 9 respectively. Notably, the feature set was augmented using BLP. In both datasets, SVM outperformed the other learners. Therefore, we selected it as the meta learner of our stacking ensemble.

**Table 8:** 10-Fold CV performance of 10 ML models on N-GlycositeAtlas-TS-40, balanced with random undersampling. The feature set was augmented using BLP.

| Classifier | SN | SP | BACC | ACC | PREC | F1 | MCC | AUROC | AUPR |
| --- | --- | --- | --- | --- | --- | --- | --- | --- | --- |
| SVM | **0.858** | 0.758 | **0.808** | **0.808** | 0.78 | **0.817** | **0.619** | 0.882 | 0.859 |
| XGB | 0.833 | **0.775** | 0.804 | 0.804 | **0.788** | 0.809 | 0.609 | **0.885** | **0.868** |
| ET | 0.844 | 0.757 | 0.801 | 0.801 | 0.777 | 0.809 | 0.604 | 0.882 | 0.858 |
| RF | 0.849 | 0.757 | 0.803 | 0.803 | 0.778 | 0.812 | 0.609 | 0.882 | 0.861 |
| MLP | 0.792 | 0.775 | 0.784 | 0.784 | 0.779 | 0.785 | 0.568 | 0.865 | 0.842 |
| PLS | 0.857 | 0.676 | 0.766 | 0.766 | 0.726 | 0.786 | 0.542 | 0.837 | 0.796 |
| KNN | 0.757 | 0.733 | 0.745 | 0.745 | 0.739 | 0.748 | 0.49 | 0.812 | 0.767 |
| LR | 0.719 | 0.73 | 0.725 | 0.725 | 0.727 | 0.723 | 0.45 | 0.798 | 0.77 |
| NB | 0.719 | 0.708 | 0.713 | 0.713 | 0.711 | 0.715 | 0.427 | 0.78 | 0.708 |
| DT | 0.72 | 0.715 | 0.718 | 0.718 | 0.716 | 0.718 | 0.435 | 0.718 | 0.656 |

**Table 9:** 10-Fold CV performance of 10 ML models on N-GlyDE-TS-40, balanced with random undersampling.

| Classifier | SN | SP | BACC | ACC | PREC | F1 | MCC | AUROC | AUPR |
| --- | --- | --- | --- | --- | --- | --- | --- | --- | --- |
| SVM | 0.946 | 0.765 | **0.856** | **0.856** | 0.801 | **0.867** | **0.724** | **0.923** | **0.909** |
| XGB | 0.83 | 0.825 | 0.828 | 0.828 | 0.828 | 0.828 | 0.657 | 0.886 | 0.842 |
| ET | 0.956 | 0.718 | 0.837 | 0.837 | 0.773 | 0.855 | 0.695 | 0.889 | 0.846 |
| MLP | 0.864 | **0.828** | 0.846 | 0.846 | **0.835** | 0.849 | 0.693 | 0.911 | 0.887 |
| PLS | 0.932 | 0.75 | 0.841 | 0.841 | 0.789 | 0.854 | 0.694 | 0.898 | 0.871 |
| LR | 0.852 | 0.808 | 0.83 | 0.83 | 0.818 | 0.834 | 0.662 | 0.901 | 0.881 |
| NB | **0.959** | 0.711 | 0.835 | 0.835 | 0.769 | 0.853 | 0.692 | 0.845 | 0.768 |
| RF | 0.825 | 0.767 | 0.796 | 0.796 | 0.782 | 0.801 | 0.597 | 0.849 | 0.795 |
| DT | 0.808 | 0.793 | 0.801 | 0.801 | 0.801 | 0.803 | 0.604 | 0.801 | 0.744 |
| KNN | 0.871 | 0.775 | 0.823 | 0.823 | 0.797 | 0.831 | 0.651 | 0.879 | 0.826 |

### 3.3 Hyperparameter tuning

We used Grid Search to tune the hyperparameters of the different classifiers. The searched grids and the finally selected values for each classifier are reported in Table 10. The full table of each learner with their performances at the different parameter combinations is given in Supplementary.

**Table 10:**
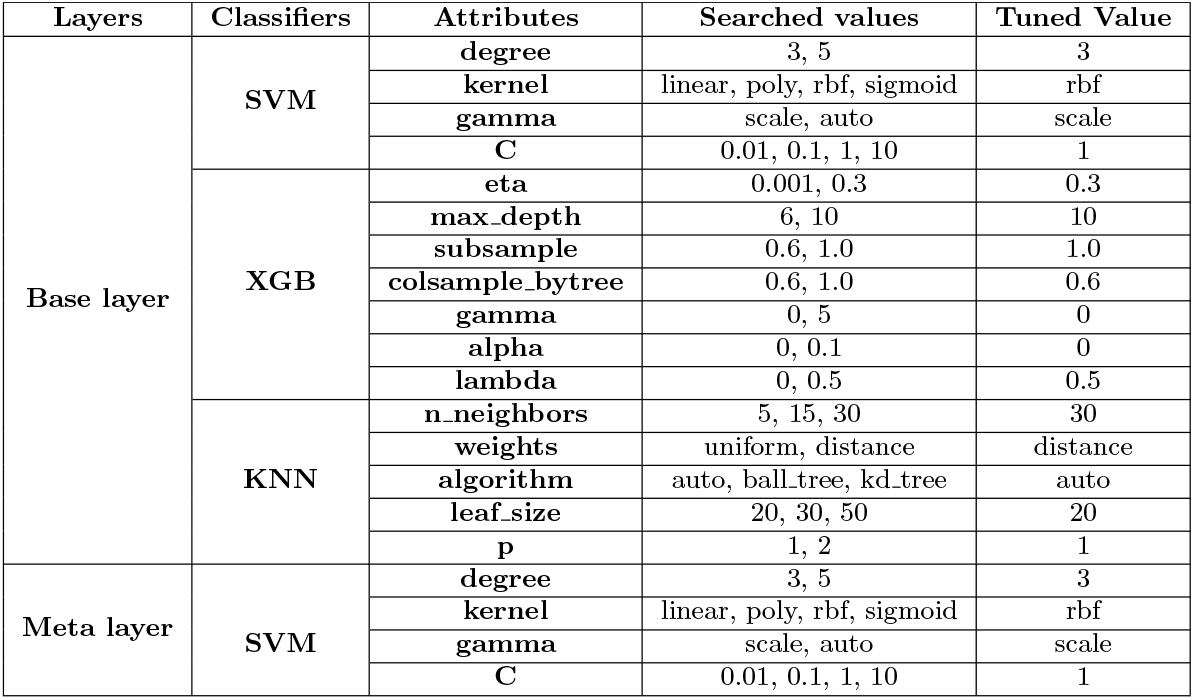
List of learner hyperparameters and their final (tuned) values.

**Table 11:** Feature combinations.

| Feature combination label | Feature set |
| --- | --- |
| FC-1 | ProteinBERT, ESM-2-Window, ProtT5-XL-U50-Per-Residue |
| FC-2 | ProteinBERT, ESM-2-Window, ProtT5-XL-U50-Per-Residue, BLP |

### 3.4 Significance of BLP on the performance

To examine the benefit of the base learner probabilities (BLP) being added to the feature vector before it is passed through the meta layer of the stacking ensemble, we compared the performance of the meta classifier without and with the BLP. These two variations of feature vectors are referred to as FC-1 and FC-2 respectively (Table 11). The results are shown in Figures 3 and 4. In both cases, we can see that FC-2 performs better than FC-1 in terms of almost all metrics. Thus the contribution of BLP in improving the efficacy of the model is evident.

**Figure 3.**
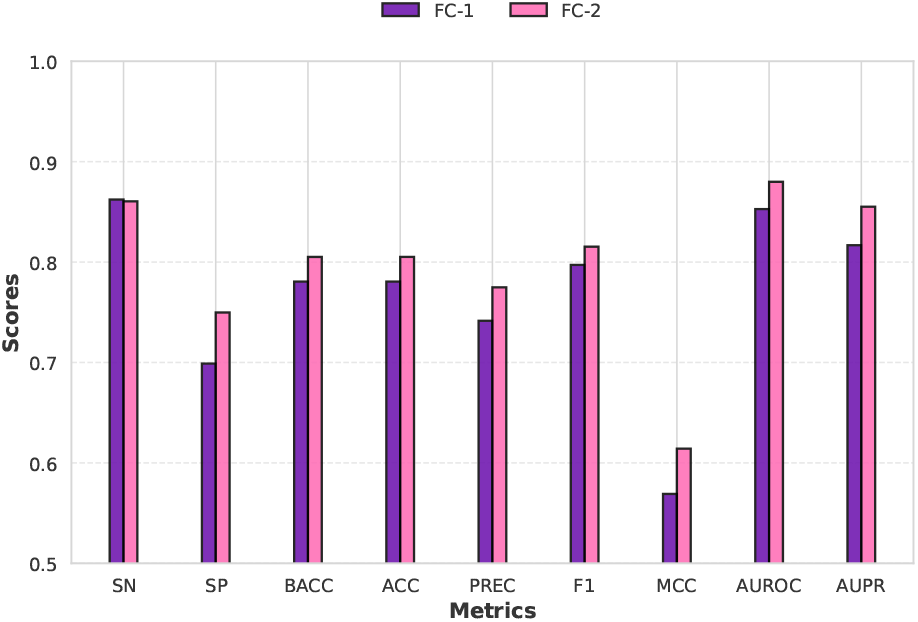
10-Fold CV performance of meta classifier without BLP (FC-1) and with BLP (FC-2) on the randomly undersampled N-GlycositeAtlas-TS-40 set.

**Figure 4.**
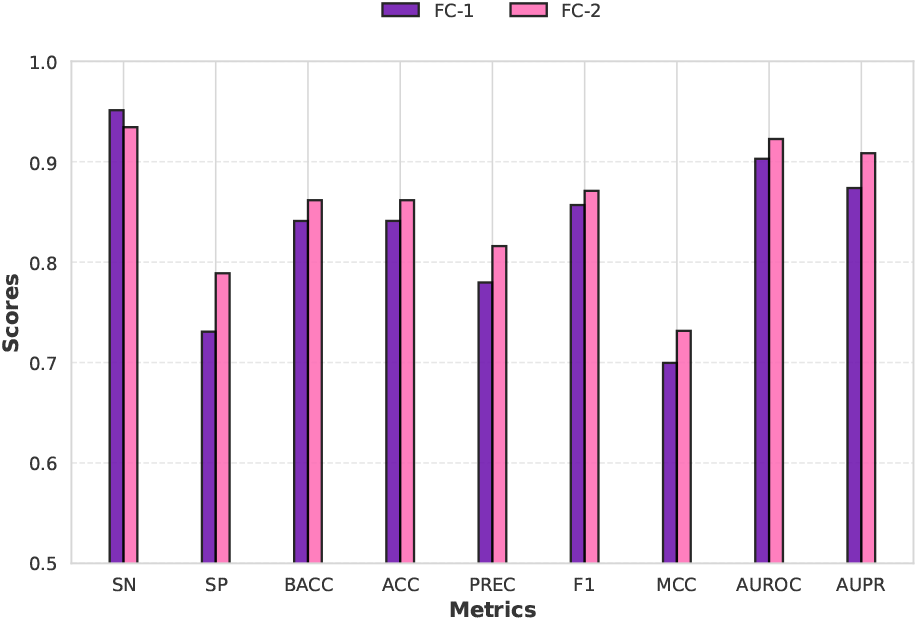
10-Fold CV performance of meta classifier without BLP (FC-1) and with BLP (FC-2) on the randomly undersampled N-GlyDE-TS-40 set.

To further examine the contribution of BLP as well as the other feature groups, we performed SHAP [32] analysis as follows. The dimension of our feature vector is 2846 with BLP (1024 from ProtT5-XL-U50-Per-Residue, 1280 from ESM-2-Window, 512 from ProteinBERT and 30 from BLP). As it is prohibitively expensive to conduct SHAP analysis with large feature vectors, we reduced the feature dimension to 100 using PCA (Principal component analysis) [45]. We applied PCA independently on each feature group and collected 10 principal components (PCs) from the BLP vector and 30 PCs each from the other three feature groups. These were then concatenated to derive the 100-dimensional reduced feature space. As the computation time of SHAP also increases considerably with sample size, we used sampling in the N-GlycositeAtlas-TS-40 dataset to choose 1078 samples, comprising equal number of positive and negative samples. As the N-GlyDE-TS-40 dataset was relatively small, we only used random undersampling to balance it, which resulted in a total of 824 samples. SHAP analysis was then performed and SHAP values were obtained for each feature of each sample. Using these, the mean absolute SHAP values were computed for each feature group, which are shown in the bar plots of Figures 5 (for N-GlycositeAtlas-TS-40 dataset) and 6 (for N-GlyDE-TS-40 dataset). In both cases, BLP has a considerably higher mean absolute SHAP value compared to the other feature groups. This indicates the efficacy of BLP in discriminating between the positive and negative classes.

**Figure 5.**
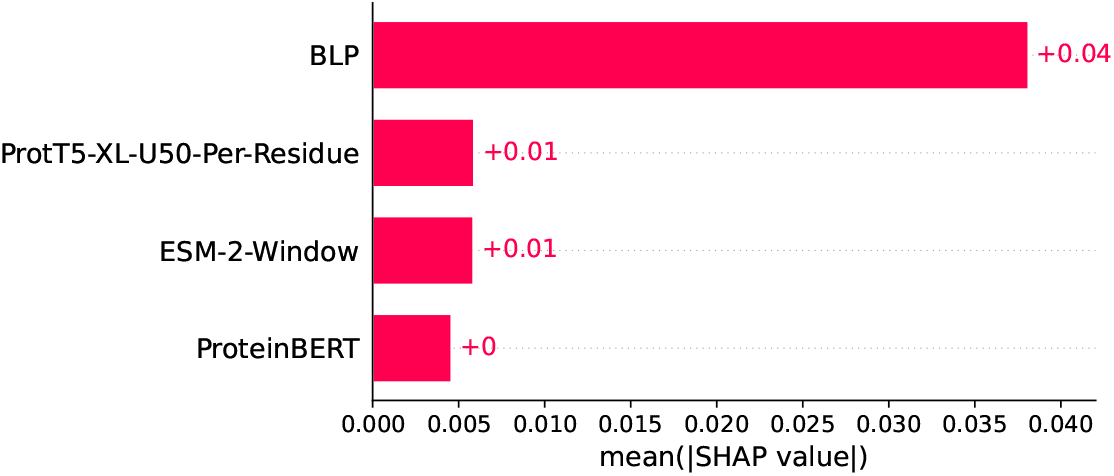
Bar plot of mean absolute SHAP values of each feature group across samples of N-GlycositeAtlas-TS-40 dataset.

**Figure 6.**
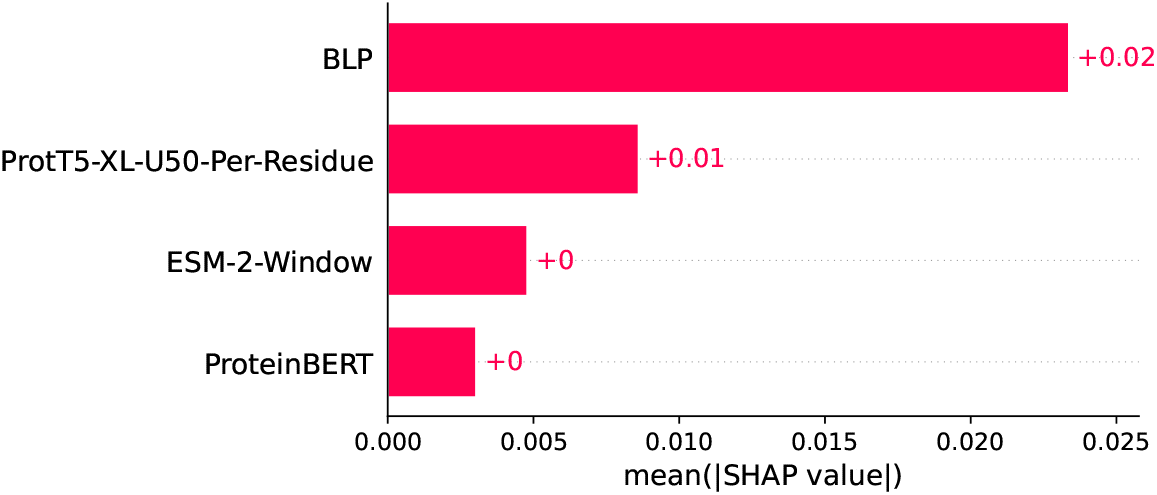
Bar plot of mean absolute SHAP values of each feature group across samples of N-GlyDE-TS-40 dataset.

### 3.5 Comparison with the state-of-the-art methods

Performance of different state-of-the-art (SOTA) methods has been compared with StackGlyEmbed on the N-GlycositeAtlas-IT and the N-GlyDE-IT datasets respectively in Tables 12 and 13. In both cases, StackGlyEmbed outperformed SOTA methods in terms of SN, F1, MCC, AUROC and AUPR. When trained on the N-GlycositeAtlas-TS dataset, it has 8.63% more sensitivity, 2.06% more F1-score, and 3.83% more MCC than LMNglyPred on the N-GlycositeAtlas-IT dataset. On the other hand, StackGlyEmbed trained on N-GlyDE-TS has 31.46% more sensitivity, 10% more F1-score, and 15.04% more MCC than LMNglyPred, and has 0.72% more sensitivity and 12.23% more MCC than EMNGly on N-GlyDE-IT dataset. Our model significantly increased the sensitivity on the N-GlyDE-IT dataset compared to the SOTA methods.

**Table 12:** Comparison of StackGlyEmbed with SOTA methods on the N-GlycositeAtlas-IT dataset.

| Predictor | SN | SP | BACC | ACC | PREC | F1 | MCC | AUROC | AUPR |
| --- | --- | --- | --- | --- | --- | --- | --- | --- | --- |
| StackGlyEmbed | **0.831** | 0.714 | **0.772** | 0.753 | 0.594 | **0.693** | **0.515** | **0.836** | **0.659** |
| LMNglyPred | 0.765 | **0.754** | 0.759 | **0.757** | **0.61** | 0.679 | 0.496 | 0.759 | 0.545 |

**Table 13:** Comparison of StackGlyEmbed with SOTA methods on the N-GlyDE-IT dataset.

| Predictor | SN | SP | BACC | ACC | PREC | F1 | MCC | AUROC | AUPR |
| --- | --- | --- | --- | --- | --- | --- | --- | --- | --- |
| StackGlyEmbed | **0.982** | 0.868 | **0.925** | **0.91** | 0.815 | **0.891** | **0.826** | **0.967** | **0.931** |
| LMNglyPred | 0.747 | **0.942** | 0.845 | 0.869 | **0.886** | 0.81 | 0.718 | 0.845 | 0.756 |
| DeepNGlyPred | 0.725 | 0.811 | 0.768 | 0.779 | 0.695 | 0.71 | 0.531 | 0.768 | 0.607 |
| EMNGly<sup>a</sup> | 0.975 | 0.7 | - | 0.884 | - | - | 0.736 | 0.946 | - |
| N-GlyDE<sup>b</sup> | 0.826 | 0.689 | - | 0.74 | 0.613 | - | 0.499 | - | - |
| GlycoMine<sup>b</sup> | 0.70 | 0.739 | - | 0.725 | 0.616 | - | 0.43 | - | - |
| NetNGlyc<sup>b</sup> | 0.844 | 0.411 | - | 0.572 | 0.46 | - | 0.265 | - | - |
| GlycoEP_Std_PPP<sup>b</sup> | 0.512 | 0.61 | - | 0.574 | 0.437 | - | 0.119 | - | - |

<sup>a</sup>Taken from [25], <sup>b</sup>Taken from [23]

For the N-GlycositeAtlas dataset, we could only consider LMNglyPred as the comparative method. There are two other SOTA methods, namely DeepNGlyPred and EMNGly, which have also utilized the N-GlycositeAtlas dataset. However, DeepNGlyPred used the entire N-GlycositeAtlas dataset to train their model. Therefore, N-GlycositeAtlas-IT cannot be used to independently assess its performance for a comparative analysis. EMNGly, on the other hand, augmented the N-GlycositeAtlas dataset (both the training and testing parts) with data from several other sources. Therefore, performance reports from their paper cannot be used for comparison. Instead, we have to use the EMNGly model for inference on N-GlycositeAtlas-IT and calculate the various metrics. However, the GitHub repository of EMNGly did not contain the required features for training and making predictions. We contacted the authors via email but could not obtain the required feature CSV files.

The ROC (Receiver Characteristic curve) curves of StackGlyEmbed and LMNglyPred in N-GlycositeAtlas, and StackGlyEmbed, LMNglyPred and DeepNGlyPred in N-GlyDE are plotted in Figure 7. Our model significantly performs better than the other existing models. Notably, the models perform worse when trained on N-GlycositeAtlas-TS vs. on N-GlyDE-TS. The authors of LMNglyPred reasoned that the N-GlycositeAtlas dataset is not properly validated and a probable cause is the possibility of spontaneous deamidation [46]. Accordingly, StackGlyEmbed trained on the N-GlyDE-TS dataset has been determined as our final prediction model and our GitHub scripts use this model for inference tasks.

**Figure 7.**
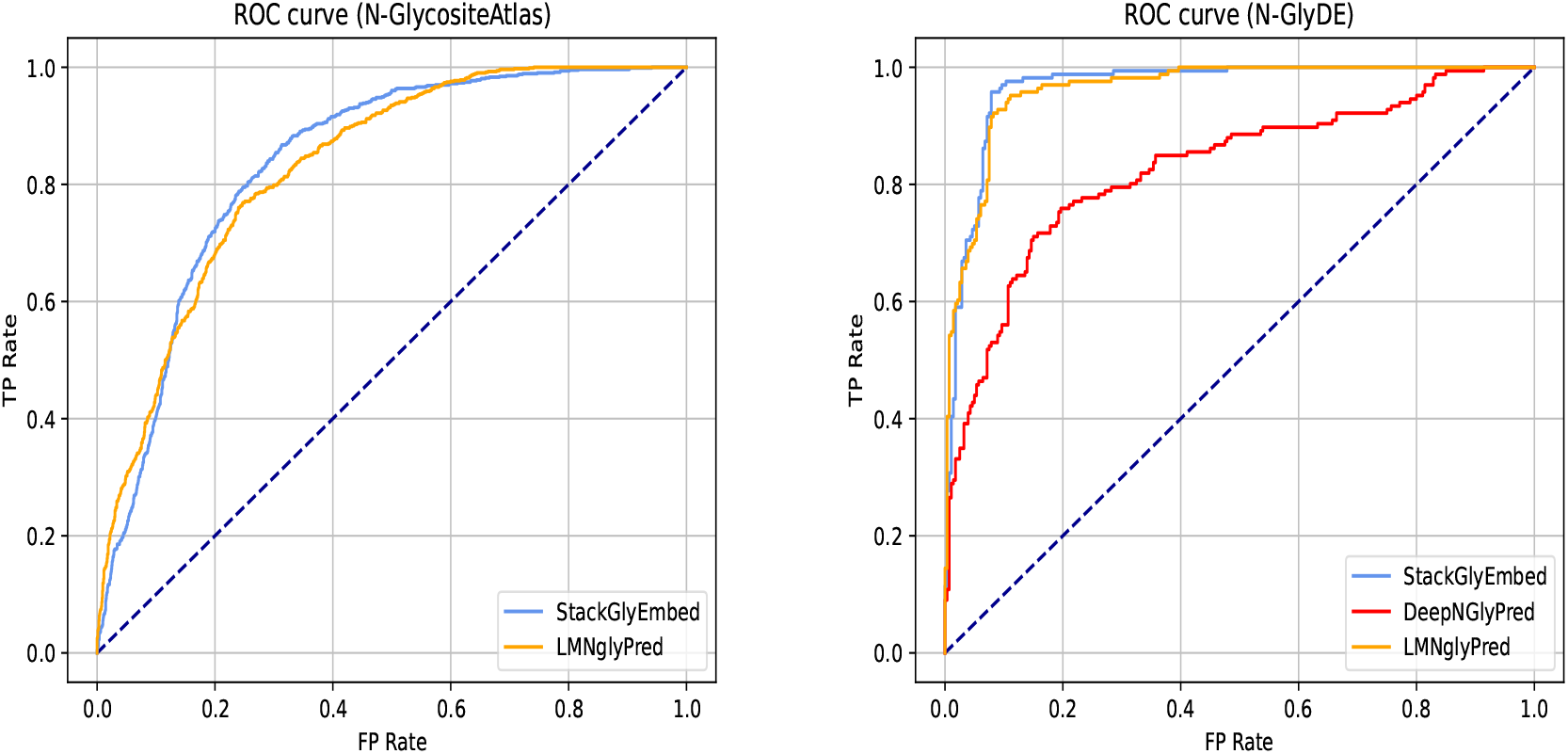
Comparison of ROC curves of StackGlyEmbed and SOTA methods on the independent sets with models trained on the corresponding training sets.

## 4 Discussion and Conclusion

In this paper, we have proposed StackGlyEmbed, a novel prediction model for N-linked glycosylation sites. We trained and tested StackGlyEmbed on two datasets, N-GlycositeAtlas and N-GlyDE. In determining a suitable representation of the residues, we have explored embeddings from several protein language models (ProtT5-XL-U50, ProteinBERT and ESM2) as well as physicochemical properties. We experimented with per-residue as well as window embeddings. As the datasets were imbalanced, we used an ensemble random undersampling approach for the base layer training with the training dataset. For the meta layer training, we used the validation set balanced using random undersampling. The base layer learners (SVM, XGB and KNN) were chosen using incremental mutual information (IMI) analysis while SVM was chosen as meta learner based on the 10-Fold CV analysis on the validation set. The fact that the same optimal feature set and the same set of base learners were selected in both datasets, is a testament to the robustness of our methodology and generalizability of the produced predictor. Indeed StackGlyEmbed showed consistent and superior performance in both independent test sets, achieving 83.1% sensitivity, 69.3% F1-score, 51.5% MCC and 65.9% AUPR on N-GlycositeAtlas-IT dataset, and 98.2% sensitivity, 89.1% F1-score, 82.6% MCC, and 93.1% AUPR on the N-GlyDE-IT dataset.

Although ensemble models are a well studied in bioinformatics, our work demonstrates the first application of stacking framework in N-linked glycosylation prediction tasks confined to N-X-[S/T] sequons. The efficacy of stacking has been demonstrated through SHAP analysis, establishing base learner probabilities (BLP) as the most important feature group. Another novelty of our work is that we have considered a windowing approach during featurization, while most studies restricted to N-X-[S/T] sequon have not. Windowing gives better localization as it considers the neighboring residues that may play a significant role in the glycosylation bindings. One of the final feature groups (ESM-2-Window) in our predictor utilizes windowing to have a positive impact on the prediction performance.

While researchers these days rush to build deep learning (DL) models for any prediction tasks, our results suggest that traditional ML models should not be ruled out. As a matter of fact, DL methods have been outperformed by traditional ML models in numerous recent bioinformatics studies [47–51]. Along that line, StackGlyEmbed too out-performed SOTA methods, many of which were based on deep learning architectures (LMNglyPred, DeepNGlyPred, NetNGlyc). Additionally, we also trained a feed-forward neural network (FNN) with our optimal set of feature groups on the N-GlycositeAtlas-TS and N-GlyDE-TS sets and respectively tested on N-GlycositeAtlas-IT and N-GlyDE-IT. The FNN delivered subpar performance compared to StackGlyEmbed, achieving 0.751 BACC, 0.669 F1, and 0.482 MCC on the N-GlycositeAtlas-IS set, and 0.894 BACC, 0.866 F1, and 0.785 MCC on the N-GlyDE-IS set. Deep learning models, including FNNs, typically require a significant amount of data to generalize well. The inferior performance of this FNN as well as SOTA DL methods could be due to the fact that DL models do not perform well under small datasets as they tend to have many parameters to learn, resulting in overfitting [52].

Our work is not without limitations. The SHAP analysis had to be conducted on a reduced feature space to fit the computational requirements within our resources. A better, more informative analysis can be done on the actual feature space in future, given a considerable amount of computing resources. Another limitation is that our model does not consider the 3d structure of proteins. While it can produce improved prediction performance, it can also limit its applicability as novel proteins are sequenced at a much faster rate than their 3d structures are experimentally determined. To solve this quandary, predicted 3d structures from AlphaFold [53] or alike could be used. However, the high computational demands and resource-intensive nature of such models prevented us from using predicted PDBs. For example, the generation of a predicted PDB file for a protein sequence with 1280 residues required ≈ 4 hours with 16 GB of memory, and 15 GB of VRAM on an Intel(R) Xeon(R) CPU @ 2.20GHz and a Tesla T4 GPU. We recognize the potential of using structural features from predicted PDBs and leave that for future research.

StackGlyEmbed brings a substantial enhancement in the N-linked glycosylation site prediction task. It is freely available as open-source scripts at https://github.com/nafcoder/StackGlyEmbed. The prediction of glycosylation sites in novel proteins can be guided by this novel predictor. New glycosylation sites and their functional implications in biological systems may be discovered faster as a result. We genuinely hope that our easy-to-use and light-weight predictor will greatly aid in predicting N-linked glycosylation sites, thereby contribute in downstream tasks that depend on it, such as, protein folding and stability, biological functions, immune responses, cell-to-cell communication, etc.

## 5 Declarations

### Availability of data and materials

Datasets, StackGlyEmbed model and scripts to reproduce the results are available at https://github.com/nafcoder/StackGlyEmbed.

### Conflict of interest

The authors declare that they have no conflict of interest.

### Author contributions

MMIN conceived the study. MMIN conducted the experiments and wrote scripts. MSR and MMIN analyzed the experimental results. All the authors reviewed the manuscript.

## Acknowledgements

MMIN is supported by the RISE Student Research Grant [No. S2024-01-004], administered by RISE, BUET. MSR is partially supported by Basic Research Grant administered by BUET.

## Notes

https://github.com/nafcoder/StackGlyEmbed

### Competing Interest Statement

The authors have declared no competing interest.

